# Interpretable dimensionality reduction of single cell transcriptome data with deep generative models

**DOI:** 10.1101/178624

**Authors:** Jiarui Ding, Anne Condon, Sohrab P. Shah

## Abstract

Single-cell RNA-sequencing has great potential to discover cell types, identify cell states, trace development lineages, and reconstruct the spatial organization of cells. However, dimension reduction to interpret structure in single-cell sequencing data remains a challenge. Existing algorithms are either not able to uncover the clustering structures in the data, or lose global information such as groups of clusters that are close to each other. We present a robust statistical model, scvis, to capture and visualize the low-dimensional structures in single-cell gene expression data. Simulation results demonstrate that low-dimensional representations learned by scvis preserve both the local and global neighbour structures in the data. In addition, scvis is robust to the number of data points and learns a probabilistic parametric mapping function to add new data points to an existing embedding. We then use scvis to analyze four single-cell RNA-sequencing datasets, exemplifying interpretable two-dimensional representations of the high-dimensional single-cell RNA-sequencing data.

## Background

Categorizing cell types comprising a specific organ or disease tissue is critical for comprehensive study of tissue development and function^1^. For example, in cancer, identifying constituent cell types in the tumor microenvironment together with malignant cell populations will improve understanding of cancer initialization, progression, and treatment response^2,3^. Technical developments have made it possible to measure the DNA and/or RNA molecules in single cells by single-cell sequencing^4–15^, or protein content by flow or mass cytometry^16,17^. The data generated by these technologies enable us to quantify cell types, identify cell states, trace development lineages, and reconstruct the spatial organization of cells^18, 19^. An unsolved challenge is to develop robust computational methods to analyze large-scale single cell data measuring the expression of dozens of protein markers to all the mRNA expressions in tens of thousands to millions of cells in order to distill single cell biology^20–23^.

Single-cell datasets are typically high-dimensional in large numbers of measured cells. For example, single-cell RNA sequencing (scRNA-seq)^19,24–26^ can theoretically measure the expression of all the genes in tens of thousands of cells in a single experiment^9,10,14,15^.For analysis, dimensionality reduction projecting high-dimensional data into low dimensional space (typically two or three dimensions) to visualize the cluster structures^27–29^ and development trajectories^30–33^ is commonly used. Linear projection methods such as principal component analysis (PCA) typically cannot represent the complex structures of single cell data in low dimensional spaces. Nonlinear dimension reduction, such as the t-distributed stochastic neighbour embedding algorithm (t-SNE)^34–39^, has shown reasonable results for many applications and has been widely used in single-cell data processing^1,40,41^. However, t-SNE has several limitations^42^. First, unlike PCA, it is a non-parametric method that does not learn a parametric mapping. Therefore, it is not natural to add new data to an existing t-SNE embedding. Instead, we typically need to combine all the data together and rerun t-SNE. Second, as a non-parametric method, the algorithm is sensitive to hyperparameter settings. Third, t-SNE is not scalable to large datasets because it has a time complexity of *O*(*N*^2^*D*) and space complexity of *O*(*N*^2^), where N is the number of cells, and D is the number of expressed genes in the case of scRNA-seq data. Fourth, t-SNE only outputs the low-dimensional coordinates but without any uncertainties of the embedding. Finally, t-SNE typically preserves the local clustering structures very well given proper hyperparameters, but more global structures such as a group of sub-clusters that forms a big cluster are missed in the low-dimensional embedding.

In this paper, we introduce a robust latent variable model, scvis to capture underlying low-dimensional structures in scRNA-seq data. As a probabilistic generative model, our method learns a parametric mapping from the high-dimensional space to a low-dimensional embedding. Therefore, new data points can be directly added to an existing embedding by the mapping function. Moreover, scvis estimates the uncertainty of mapping a highdimensional point to a low-dimensional space which adds rich capacity to interpret results. We show that scvis has superior distance preserving properties in its low-dimensional projections leading to robust identification of cell types in the presence of noise or ambiguous measurements. We extensively tested our method on simulated data and several scRNA-seq datasets in both normal and malignant tissues to demonstrate the robustness of our method.

## Results

### Modelling and visualizing single-cell RNA-sequencing data

Although scRNA-seq dataset have high dimensionality, their intrinsic dimensionalities are typically much lower. For example, factors such as cell type and patient origin explain much of the variation in a study of metastatic melanoma^3^. We therefore assume that for a high-dimensional scRNA-seq dataset 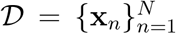 with *N* cells, where **x***_n_* is the expression vector of cell *n*,the **x***_n_* distribution is governed by a latent low-dimensional random vector **z***_n_*.For visualization purposes, the dimensionality *d* of **z***_n_* is typically two or three. We assume that **z***_n_* is distributed according to a prior, with the joint posterior distribution of the whole model as *p*(**z***_n_* | ***θ***)*p*(**x***_n_* | **z***_n_*, ***θ***). For simplicity, we can choose a factorized standard normal distribution for the prior 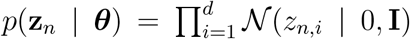. The distribution 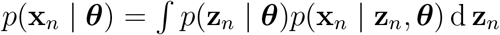 can be a complex multimodal high-dimensional distribution. To represent complex high-dimensional distributions, we assume that *p*(**x***_n_* | **z***_n_*, ***θ***) is a location-scale family distribution with location parameter ***μ**_θ_* (**z***_n_*) and scale parameter ***σ****_θ_* (**z***_n_*); both are functions of **z***_n_* parameterized by a neural network with parameter ***θ***. The inference problem is to compute the posterior distribution *p*(**z***_n_* | **x***_n_*, ***θ***), which is however intractable to compute. We therefore use a variational distribution *q*(**z***_n_* | **x***_n_*, *ϕ*) to approximate the posterior. Here *q*(**z***_n_* | **x***_n_*, *ϕ*) is a multivariate normal distribution with mean ***μ**_ϕ_* (**x***_n_*) and standard deviation ***σ****_ϕ_* (**x***_n_*). Both parameters are (continuous) functions of **x***_n_* parameterized by a neural network with parameter *ϕ*. To model the data distribution well (with a high likelihood of 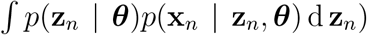, the model tends to assign similar posterior distributions *p*(**z***_n_* | **x***_n_*, ***θ***) to cells with similar expression profiles. To explicitly encourage cells with similar expression profiles to be proximal (and those with dissimilar profiles to be distal) in the latent space, we add the t-SNE objective function on the latent z distribution as a constraint. More details about the model and the inference algorithms are presented in the Methods section. The scvis model is implemented in Python using Tensorflow^43^ with a command-line interface and is freely available.

### Single-cell datasets

We analyzed four single-cell RNA sequencing (scRNA-seq) datasets in this study^1,3,9,44^. Data were mostly downloaded from the single-cell portal (https://portals.broadinstitute.org/single_cell). Two of these datasets were originally used to study intra-tumour heterogeneity and the tumour-microenvironment in metastatic melanoma^3^ and oligodendroglioma^44^, respectively. One dataset was used to categorize the mouse bipolar cell populations of the retina^1^, and one dataset was used to categorize all cell types in the mouse retina^9^. For all the scRNA-seq datasets, we used principal component analysis (as a noise-reduction preprocessing step^1,19^) to project the cells into a 100dimensional space, and used the projected coordinates in the 100-dimensional spaces as inputs to `scvis`.

### Experimental setting and implementation

The variational approximation neural network has three hidden layers (*l*_1_,*l*_2_, and *l*_3_)with 128, 64, and 32 hidden unit seach, and the model neural network has five hidden layers 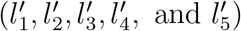 with 32, 32, 32, 64, and 128 units each. We use the exponential linear unit activation function as it has been shown to speed up the convergence of optimization^45^, and the Adam stochastic optimization algorithm with a learning rate of 0.01^46^. The time complexity to compute the t-SNE loss is quadratic in terms of the number of data points. Consequently we use mini-batch optimization and set the mini-batch size to 512 (cells). We expect that a large batch of data could be better in estimating the high-dimensional data manifold, however we found that 512 cells work accurately and efficiently in practice. We run the Adam stochastic gradient descent algorithm, for 500 epochs for each dataset with at least 3,000 iterations by default. For large datasets, running 500 epochs is computationally expensive, we therefore run the Adam algorithm for a maximum of 30,000 iteration or two epochs (which is larger). We use an L2 regularizer of 0.001 on the weights of the neural networks to prevent overfitting.

### Benchmarking against t-SNE on simulated data

To demonstrate that `scvis` can robustly learn a low-dimensional representation of the input data, we first simulated data in a two-dimensional space (for easy visualization) as in Fig. 1(a). The big cluster on the left consisted of 1,000 points and the five small clusters on the right each had 200 points. The five small clusters were very close to each other and could roughly be considered as a single big cluster. There were 200 uniformly distributed outliers around these six clusters. For each two-dimensional data point with coordinates (*x,y*), we then mapped it into a nine-dimensional space by the transformation (*x* + *y*, *x* – *y*, *xy*, *x*^2^, *y*^2^, *x*^2^ *y*, *xy*^2^, *x*^3^, *y* ^3^). Each of the nine features was then divided by its corresponding maximum absolute value.

**Figure 1:**
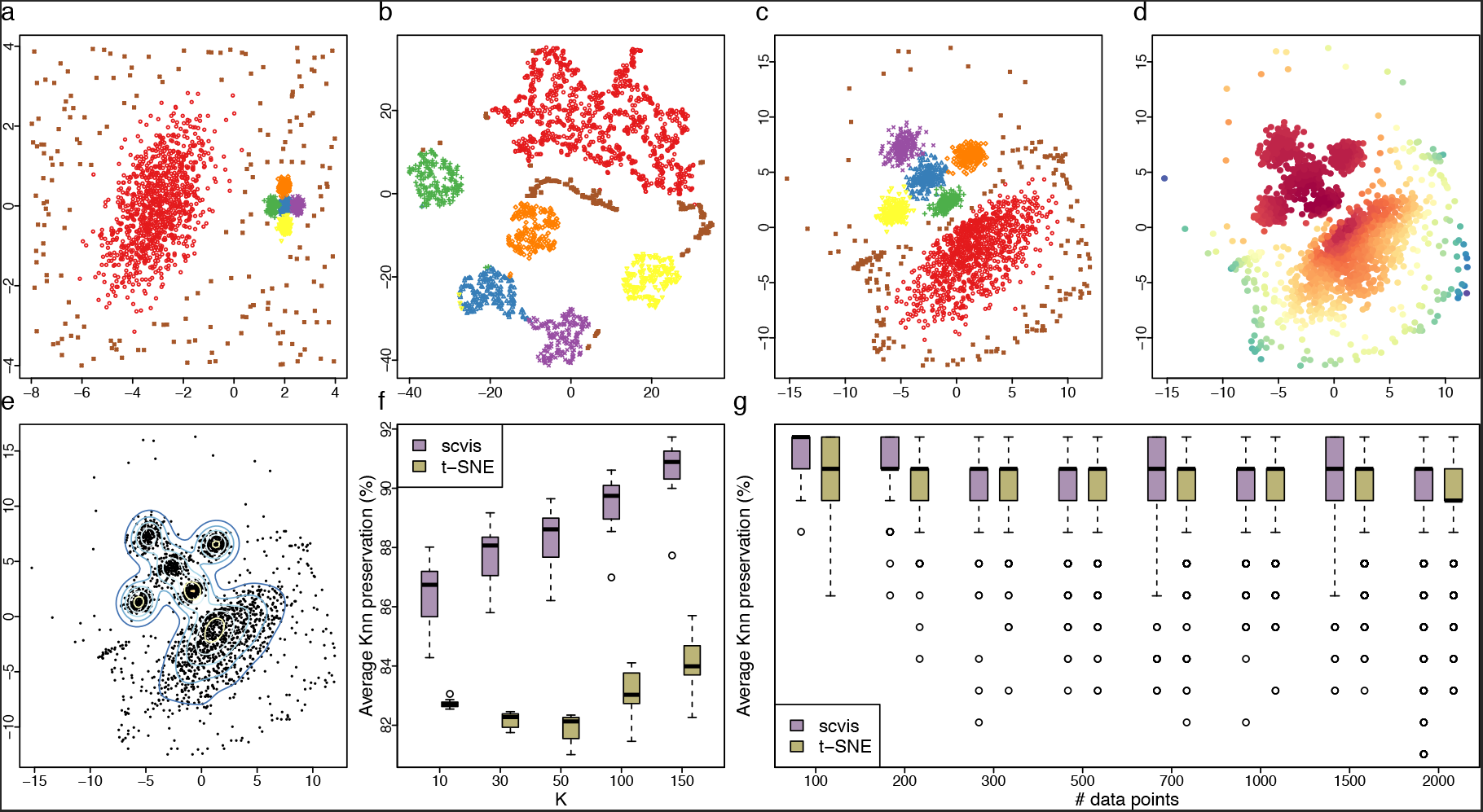
Benchmarking scvis against t-SNE on synthetic data. (a) The original 2, 200 two dimensional synthetic data points, (b) t-SNE results on the transformed nine-dimensional dataset with default perplexity parameter of 30, (c) scvis results, (d) colouring points based on their log-likelihoods from scvis, (e) the kernel density estimates of the scvis results, (f) average *K*-nearest neighbour conservations across ten runs for different *K* s, and (g) the average *K*-nearest neighbour conservations (K = 10) for different numbers of sub-sampled data points.

Although t-SNE (with default parameter setting) uncovered the six clusters in this dataset, it was still challenging to infer the overall layout of the six clusters. For example, it looked like that there were four small clusters around the big cluster, and one cluster was further away on the right (Fig. 1(b)). We could not interpret the t-SNE results this way because t-SNE by design preserves local structure of the high-dimensional data, but ‘global’ structure is not reliable. Moreover, for the uniformly distributed outliers, t-SNE put them into several compact clusters which were adjacent to other genuine clusters.

The `scvis` results, on the other hand, better preserved the overall structure of the original data (Fig. 1(c)): 1) The five small clusters were on one side, and the big cluster was on the other side. The relative positions of the clusters were also preserved, e.g., for the five small clusters, the blue cluster was in the centre and the other clusters were around the blue cluster; the green cluster was relatively closer to the big cluster than the other clusters, and the purple cluster was on the opposite side of the green cluster. The centres of the green, blue, and purple clusters formed a line, and this line passed the centre of the red cluster. 2) Outliers were scattered around the genuine cluster as in the original data. In addition, as a probabilistic generative model, `scvis` not only learned a low-dimensional representation of the input data, but also provided a way to quantify the uncertainty of the low-dimensional mapping of each input data point by its log-likelihood. For example, we coloured the lowdimensional embedding of each data point by its log-likelihood (Fig. 1(d)). We can see that generally, scvis put most of its modelling power to model the five compact clusters, while the outliers far from the five compact clusters tended to have lower log-likelihoods. Thus, by combining the log-likelihoods and the low-dimensional density information (Fig. 1(e)), we can better interpret the structure in the original data.

The low-dimensional representation may change for different runs because the scvis objective function can have different local maxima. To test the stability of the low-dimensional representations, we ran scvis ten times. Generally, the two-dimensional representations from the ten runs (Supplementary Fig. 1(a-j)) showed similar patterns as in Fig. 1(c). As a comparison, we also ran t-SNE ten times, and the results (Supplementary Fig. 1(k-t)) showed that the layouts of the clusters were less preserved, e.g., the relative positions of the clusters changed from run to run. To quantitatively compare scvis and t-SNE results, we computed the average *K*-nearest neighbour (*K* nn) preservations across runs for *K* ϵ{10, 30, 50, 100, 150}. Specifically, for the low-dimensional representation from each run, we constructed *K*nn graphs for different *K* s. We then computed the *K*nn graph from the high-dimensional data for a specific *K*. Finally, we compared the average overlap of the *K* nn graphs from the low-dimensional representations with the *K* nn graph from the highdimensional data for a specific *K*. For scvis, the median *K* nn conservations monotonically increased from 86.7% for *K* = 10, to 90.9% for *K* = 150 (Fig. 1(f)). For t-SNE, the median *K* nn conservations first decreased from 82.7% for *K* = 10, to 82.1% for *K* = 50 (because t-SNE preserves the local structures, or nearest neighbours of each point, where the number of nearest neighbours is determined by the perplexity parameter, see Methods for details), and then increased to 84.0% for *K* =150. In addition, for this dataset, scvis preserved *K* nn more effectively than t-SNE.

To test how scvis performs on smaller datasets, we sub-sampled the nine-dimensional synthetic datasets. Specifically, we sub-sampled 200, 300, 500, 700, 1,000, 1,500, and 2,000 points from the original dataset, and ran scvis on each sub-sampled dataset. We then computed the *K* nn conservations (*K* = 10), and found that the *K* nn conservations from the scvis results were significantly higher than those from t-SNE results (adjusted Wilcoxon-test p-value < 0.05 for all the sub-sampled datasets, Fig. 1(g)). scvis performs very well on all the sub-sampled datasets (Fig. 2(a-h)). Even with just 100 data points, the two-dimensional representation (Supplementary Fig. 2(a)) preserved much of the structures in the data, e.g., as for the results on the original 2,200 data points (Fig. 1(c)), the five small clusters and the big clusters were mapped to different regions. For the five small clusters, the blue cluster was in the centre and the other four clusters were around the blue cluster. The green cluster was closest to the big cluster and the purple cluster was at the opposite site of the green cluster. The log-likelihoods estimated from the sub-sampled data also recapitulated the log-likelihoods from the original 2,200 data points (Supplementary Fig. 3(a-h)). The t-SNE results on the sub-sampled datasets (Supplementary Fig. 2(i-p)), generally revealed the clustering structures. However, only for smaller datasets (e.g., 100, 200, or 300 data points), we could see that for the five small clusters, the blue cluster was in the centre, and the other four clusters were around the blue cluster. Furthermore, the relative positions of the five clusters and the big cluster were largely inaccurate. We noted that the centres of the green, blue, and purple clusters generally did not maintain the structure of the input data.

**Figure 2:**
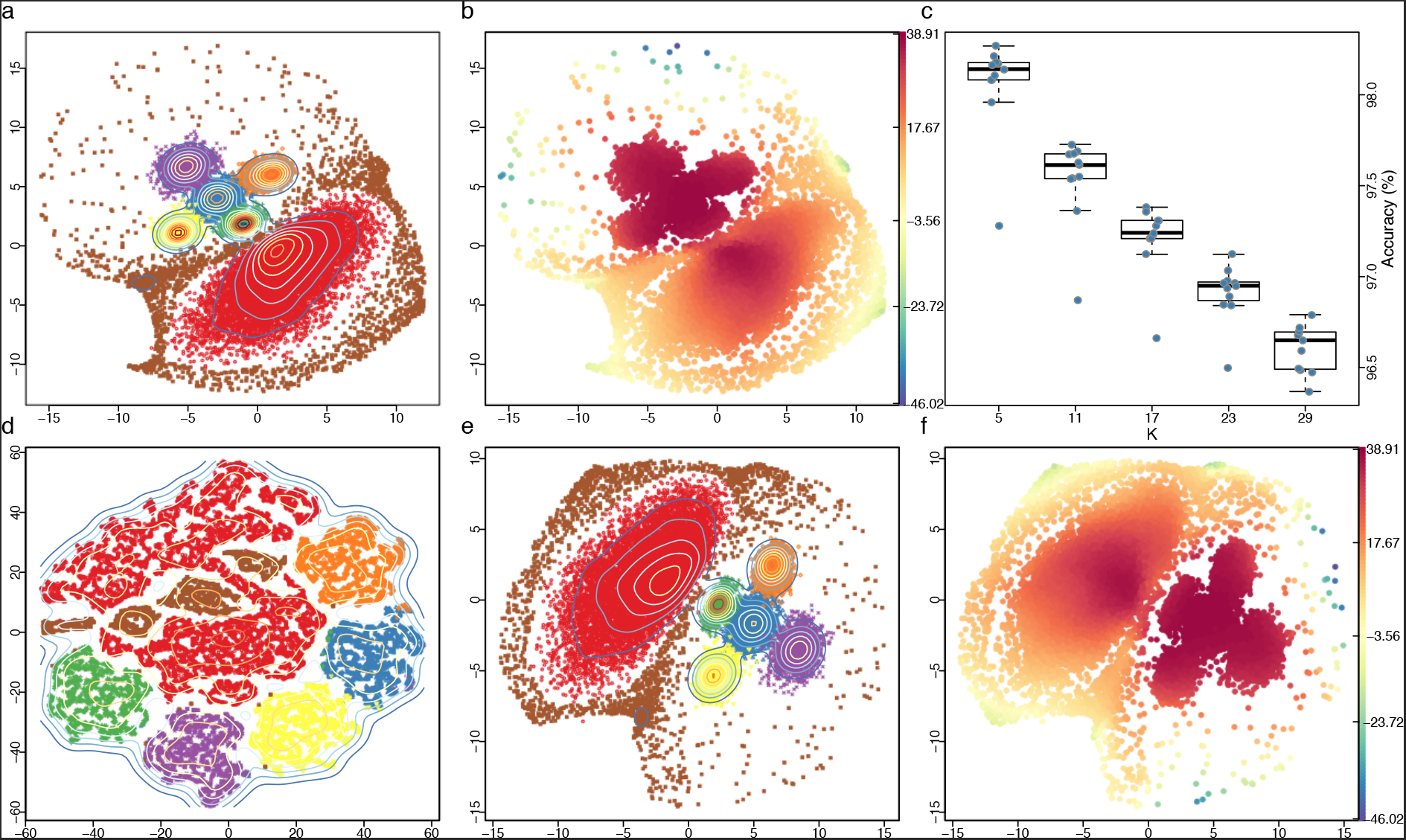
Benchmarking scvis against t-SNE on a larger synthetic dataset with 22,000 data points. (a) mapping 22, 000 new data points based on the learned probabilistic mapping function from the 2, 200 training data points, (b) the estimated log-likelihoods, (c) the average *K*-nearest neighbour classification accuracies for different *K* s across eleven runs, the classifiers were trained on the eleven embeddings as used in calculating the *K*-nearest neighbour conservations, (d) t-SNE results on the larger dataset, (e) `scvis` results on the larger dataset with the same perplexity parameter as used in Fig. 1, and (f) `scvis` log-likelihoods on the larger dataset.

**Figure 3:**
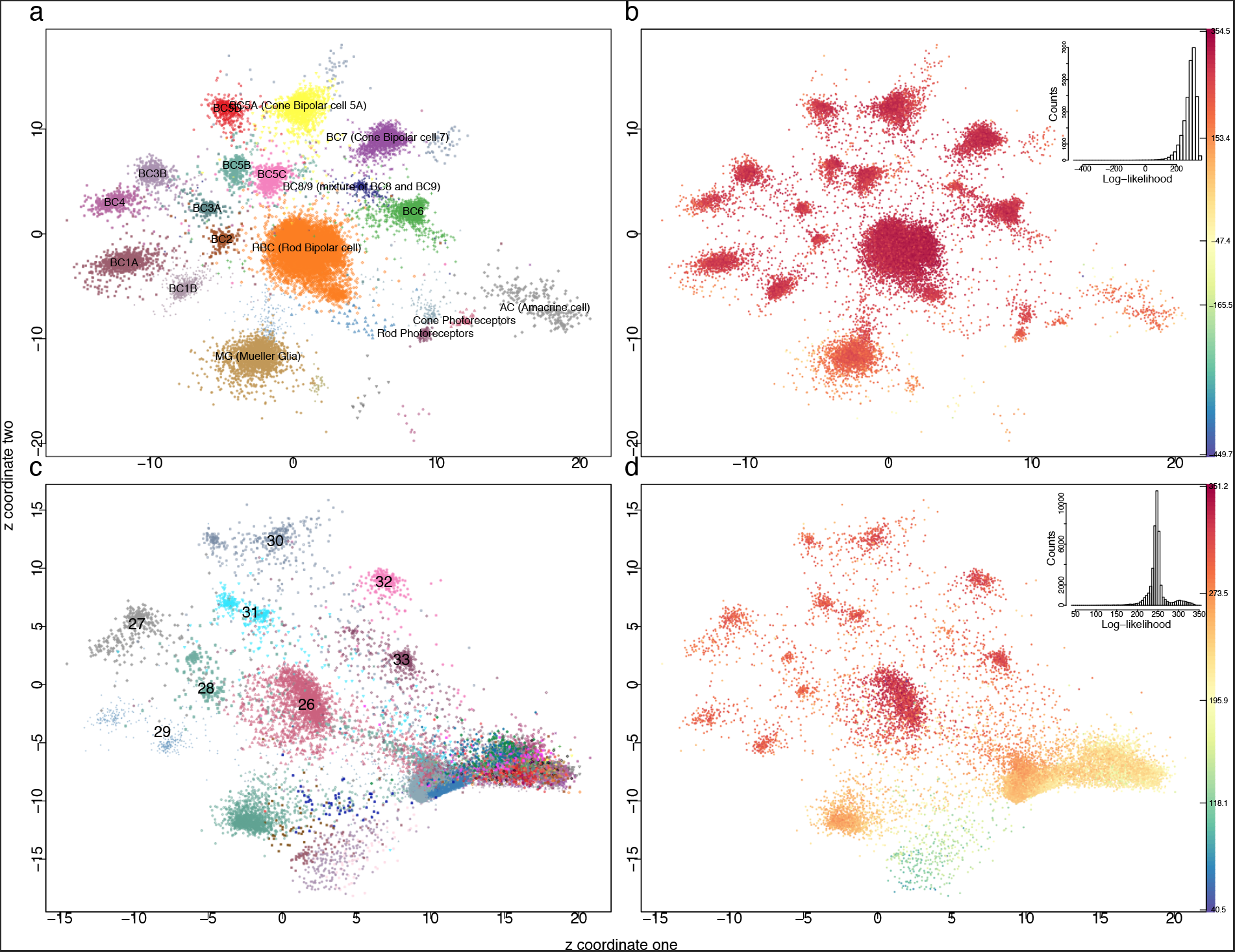
Learning a probabilistic mapping function from the bipolar data and applying the function to the independently generated mouse retina dataset. (a) scvis learned two dimensional representations of the bipolar dataset, (b) colouring each point by the estimated log-likelihood, (c) the whole mouse retina dataset was directly projected to a two dimensional space by the probabilistic mapping function learned from the bipolar data, and (d) colouring each point from the retina dataset by the estimated log-likelihoods.

To test the performance of `scvis` when adding new data to an existing embedding, we increased by tenfold the number of points in each cluster and the number of outliers (for a total of 22,000 points) using a different random seed. The embedding (Fig. 2(a-b)) was very similar to that of the 2, 200 training data points in Fig. 1(c-d). We trained K nn classifiers on the embedding of the 2, 200 training data for *K* ϵ {5, 11, 17, 23, 29}, and used the trained classifiers to classify the embedding of the 22, 000 points, repeating eleven times. Median accuracy (the proportion of points correctly assigned to their corresponding clusters) was 98.1% for *K* =5, and 96.7% for *K* =29. The performance decreased mainly because for a larger *K*, the outliers were wrongly assigned to the six genuine clusters.

As a non-parametric dimension reduction method, t-SNE was sensitive to hyperparameter setting, especially the perplexity parameter (the effective number of neighbours, see the Methods section for details). The optimal perplexity parameter increased as the total number of data points increased. In contrast, as we adopted mini-batch for training `scvis`, the perplexity parameter for scvis was stable for different numbers of input data points. For this larger dataset, the t-SNE results (Fig. 2(d)) were difficult to interpret without the ground-truth cluster information because it was already difficult to see how many clusters in this dataset, not to mention to uncover the overall structure of the data. Finally, scvis performed well on this larger dataset (Fig. 2(e-f)), without changing the perplexity parameter for `scvis`.

### Learning a parametric mapping for single-cell data

We next analyzed the scvis learned probabilistic mapping from a training single-cell dataset, and tested how it performed on unseen data. We first trained a model on the mouse bipolar cell of the retina dataset^1^, and then used the learned model to map the independently generated mouse retina dataset^9^. The two-dimensional coordinates from the bipolar dataset captured much information in this dataset (Fig. 3(a)). For example, non-bipolar cells such as Amacrine cells, Mueller Glia, and photoreceptors were at the bottom, the Rod bipolar cells were in the middle, and the cone bipolar cells were on the top left around the Rod bipolar cells. Moreover, the ‘OFF’ cone bipolar cells (BC1A, BC1B, BC2, BC3A, BC3B, BC4) were on the left and close to each other, and the ‘ON’ cone bipolar cells (BC5A-D, BC6, BC7, BC8/9) were at the top. Cell doublets and contaminants (accounting for 2.43% of the cells comprised eight clusters^1^, with distinct colour and symbol combinations in Fig. 3(a) but not labelled) were rare in the bipolar datasets, and they were mapped to low-density regions in the low-dimensional plots (Fig. 3(a)).

Consistent with the synthetic data (Fig. 1), t-SNE put the ‘outlier’ cell doublets and contaminants into very distinct compact clusters (Supplementary Fig. 4(a), t-SNE coordinates from Shekhar^1^ *et al*). In addition, although t-SNE mapped cells from different cell populations into distinct regions, more global organizations of clusters of cells were missed in the t-SNE embedding. For example, the Rod bipolar cells were mapped to the centre, and other cell clusters were around the Rod bipolar cells. The ‘ON’ cone bipolar cell clusters, the ‘OFF’ cone bipolar cell clusters, and other non-bipolar cell clusters were mixed together in the t-SNE results.

**Figure 4:**
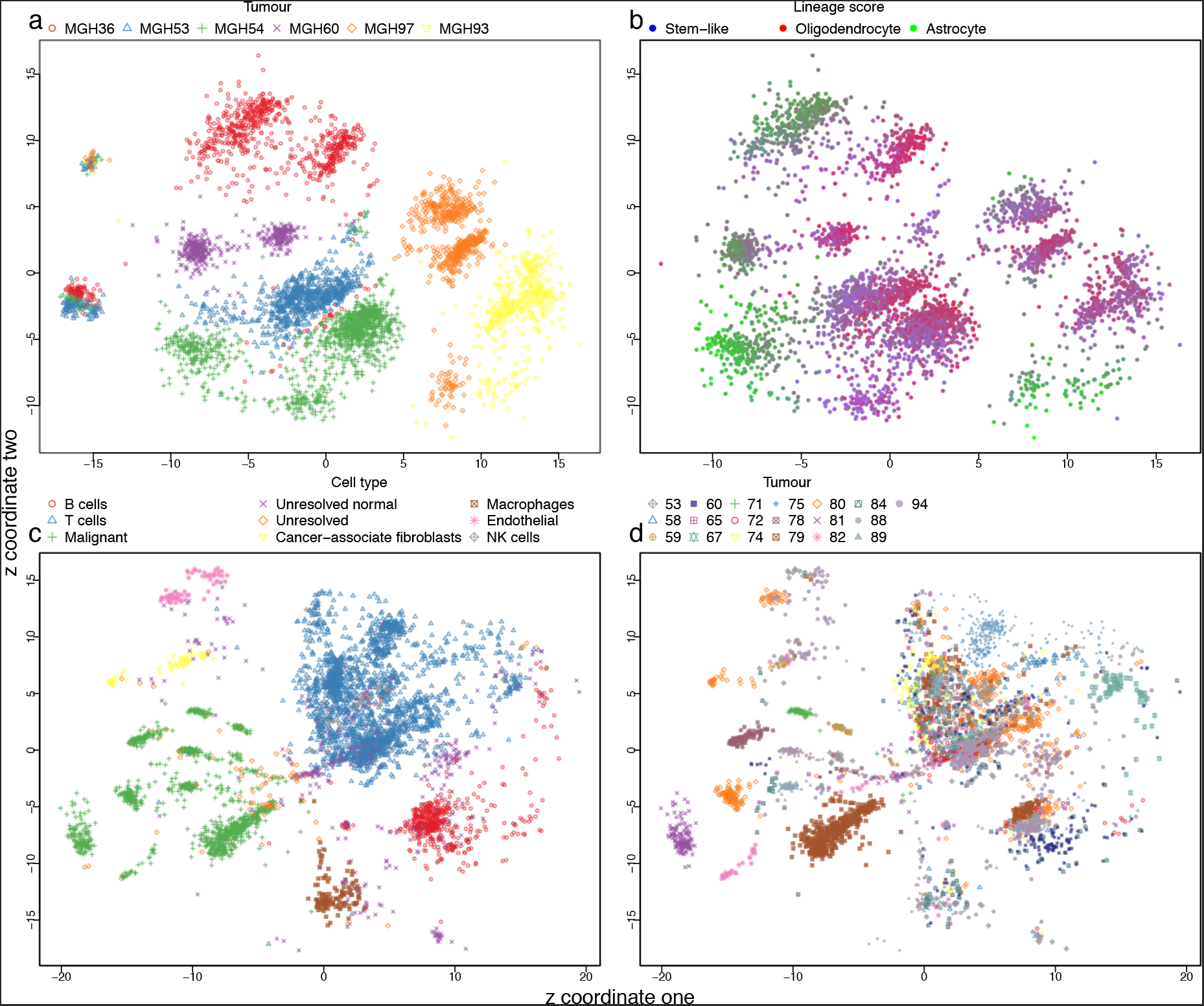
scvis learned low-dimensional representations. (a) The oligodendroglioma dataset, each cell is coloured by its patient of origin, (b) the oligodendroglioma dataset, each cell is coloured by its linage score from Tirosh *et al* ^44^, (c) the melanoma dataset, each cell is coloured by its cell type, and (d) the melanoma dataset, each cell is coloured by its patient of origin.

The bipolar cells tended to have higher log-likelihoods than non-bipolar cells such as Amacrine cells, Mueller Glia, and photoreceptors (Fig. 3(b)), suggesting the model used most of its power to model the bipolar cells, while other cell types were not modelled as well. The embedded figure at the top right corner shows the histogram of the log-likelihoods. The majority of the points exhibited high log-likelihoods (with a median of 292.4). The bipolar cells had significantly higher log-likelihoods (median log-likelihood of 298.4) relative to non-bipolar cells (including Amacrine cells, Mueller Glia, Rod and Cone photoreceptors) (median log-likelihood of 223.6; *t*-test p-value < 10^-15^; Supplementary Fig. 4(b)). The Amacrine cells had the lowest median log-likelihood (median log-likelihood for Amacrine cells, Mueller Glia, Rod and Cone photoreceptors were 226.4, 187.3, 222.7, and 205.4, respectively; Supplementary Fig. 4(b)).

We used the learned probabilistic mapping from the bipolar cells to map the independent whole retina dataset^9^. We first projected the retina dataset to the subspace spanned by the first 100 principal direction vectors of the bipolar dataset, and then mapped each 100dimensional vector to a two-dimensional space based on the learned scvis model from the bipolar dataset. The bipolar cell clusters in the retina dataset identified in the original study^9^ (cluster 26-33) tended to be mapped to the corresponding bipolar cell subtype regions discovered in the study^1^ (Fig. 3(c)). Although Macosko^9^ *et al* only identified eight subtypes of bipolar cells, all the recently identified 14 subtypes of bipolar cells^1^ were possibly present in the retina dataset as can be seen from Fig. 3(c), i.e., cluster 27 (BC3B and BC4), cluster 28 (BC2 and BC3A), cluster 29 (BC1A and BC1B), cluster 30 (BC5A and BC5D), cluster 31 (BC5B and BC5C), and cluster 33 (BC6 and BC8/9).

Interestingly, there was a cluster just above the Rod photoreceptors (Fig. 3(c)) consisting of different subtypes of bipolar cells. In the bipolar dataset, cell doublets or contaminants were mapped to this region (Fig. 3(a)). We used densitycut^47^ to cluster the two-dimensional mapping of all the bipolar cells from the retina dataset to detect this mixture of bipolar cell cluster (Supplementary Fig. 4(c), where the 1, 535 high-density points in this cluster were labeled with red circles). To test whether this mixture cell population were artifacts of the projection, we randomly drew the same number of data points from each bipolar subtype as in the mixture cluster and computed the *K*-nearest neighbours of each data point (here *K* was set to log_2_ (1535) = 11). We found that the 11-nearest neighbours of the points from the mixture clusters were also mostly from the mixture cluster (median of 11 and mean of 10.8), while for the randomly selected points from the bipolar cells, a relatively small number of points of their 11-nearest neighbours (median of 0 and mean of 0.2) were from the mixture cluster. The results suggest that the bipolar cells in the mixture cluster were substantially different from other bipolar cells. Finally, this mixture of bipolar cells had significantly lower log-likelihoods compared with other bipolar cells (t-test p-value < 1e^-15^, Supplementary Fig. 4(d)).

Non-bipolar cells especially Mueller Glia cells were mapped to the corresponding regions as in the bipolar dataset (Fig. 3(c)). Photoreceptors (Rod and Cone photoreceptors accounting for 65.6% and 4.2% of all the cells from the retina^9^) were also mapped to their corresponding regions as in the bipolar dataset (Supplementary Fig. 4(e)). The Amacrine cells (consisting of 21 clusters) together with Horizontal cells and Retinal ganglion cells were mapped to the bottom right region (Fig. 3(f)); all the Amacrine cells were assigned the same label and the same colour).

As in the training bipolar data, the bipolar cells in the retina dataset also tended to have high log-likelihoods, and other cells tended to have relatively lower log-likelihoods (Fig. 3(d)). The embedded plot on the top right corner shows a bimodal distribution of the log-likelihoods. The ‘Other’ cells types (Horizontal cells, Retina ganglion cells, Microglia cells etc) that were only in the retina dataset had the lowest log-likelihoods (median log-likelihoods of 181.7, Supplementary Fig. 4(d)).

### Analyzing tumor microenvironments and intro-tumor heterogeneity

We next used scvis to analyze tumor microenvironments and intra-tumor heterogeneity. The oligodendroglioma dataset consists of mostly malignant cells (Supplementary Fig. 5(a)). We used densitycut^47^ to cluster the two-dimensional coordinates to produce 15 clusters (Supplementary Fig. 5(b)). The non-malignant cells (Microglia/Macrophage and Oligodendrocytes) formed two small clusters on the left and each consisted of cells from different patients. We therefore computed the entropy of each cluster based on the cells of origin (enclosed bar plot). As expected, the non-malignant clusters (cluster one and cluster five) had high entropies. Cluster 12 (cells mostly from MGH53 and MGH54) and cluster 14 (cells from MGH93 and MGH94) also had high entropies (Fig. 4(a)). The cells in these two clusters consisted of mostly astrocytes (Fig. 4(b); the oligodendroglioma cells could roughly be classified as oligodendrocyte, astrocyte, or stem-like cells.) Interestingly, cluster 15 had the highest entropy, and these cells had significant higher Stem-like scores (*t*-test p-value < 10^-12^). We also coloured cells by the cell-cycle scores (G1/S scores, Supplementary Fig. 5(c); G2/M scores, Supplementary Fig. 5(d)), and found that these cells also had significantly higher G1/S scores (*t*-test p-value < 10^-12^) and G2/M scores (*t*-test p-value < 10^−9^). Therefore, cluster 15 cells consisted of mostly Stem-like cells, and these cells were cycling.

Malignant cells formed distinct clusters even if they were from the same patient (Fig. 4(a)). We next coloured each malignant cell by its lineage score^44^ (Fig. 4(b)). The cells in some clusters highly expressed the astrocyte gene markers, or the oligodendrocyte gene markers. The stem-like cells tended to be rare and they could link ‘outliers’ connecting oligodendrocyte and astrocyte cells in the two-dimensional scatter plots (Fig. 4(b)). In addition, some clusters of cells consisted of mixtures of cells (e.g., both oligodendrocyte and stem-like cells), suggesting other factors such as genetic mutations and epigenetic measurements would be required to fully interpret the clustering structures in the dataset.

For the melanoma dataset, the authors profiled both malignant cells and non-malignant cells^3^. The malignant cells originated from different patients were mapped to the bottom left region (Fig. 4(c)). These malignant cells were further subdivided by the patients of origin (Fig. 4(d)). Similar to the oligodendroglioma dataset, non-malignant immune cells such as T cells, B cells, and macrophages, even from different patients, tended to be grouped together by cell types instead of patients of origin of the cells (Fig. 4(c-d)), although for some patients (e.g., 75, 58, and 67, Fig. 4(d)), their immune cells showed patient-specific bias. We did a differential expression analysis of patient 75 T cells and other patient T cells using limma^48^. Of the top 100 differentially expressed genes, most of the highly expressed genes were Ribosome genes (Supplementary Fig. 6(a)). As housing keeping genes, ideally, Ribosome genes should be highly expressed in all the T cells suggesting perhaps that batch effects are detectable between patient 75 T cells and other patient T cells. In addition, CD8A was significantly expressed higher in patient 75 T cells, suggesting that most of the patient 75 T cells were CD8 T cells.

Interestingly, as non-malignant cells, cancer-associated fibroblasts were mapped to the region adjacent to the malignant cells. The endothelial cells were just above the cancer-associated fibroblasts (Fig. 4(d)). To test whether these cells were truly more similar with the malignant cells than with immune cells, we first computed the average principal component values in each type of cells and did a hierarchical clustering analysis (Supplementary Fig. 6(b)). Generally, there were two clusters: one cluster consisted of the immune cells and the ‘Unsolved normal’ cells, while the other cluster consisted of cancer-associated fibroblasts, endothelial cells, malignant cells, and the ‘Unsolved’ cells indicating cancer-associated fibroblasts and endothelial cells were more similar to malignant cells (they had high PC1 values) than to the immune cells.

## Discussion

We have developed a novel method, scvis for modeling and reducing dimensionality of single cell gene expression data. We demonstrated that scvis can robustly preserve the structures in high-dimensional datasets, including in datasets with small numbers of data points.

Our contribution has several important implications for the field. As a probabilistic generative model, scvis provides not only the low-dimensional coordinate for a given data point, but also the log-likelihood as a measure of the quality of the embedding. The log-likelihoods could potentially be used for outlier detection, e.g., for the bipolar cells in Fig. 3(b), the log-likelihood histogram shows a long tail of data points with relatively low log-likelihoods, suggesting some outliers in this dataset (the non-bipolar cells). The log-likelihoods could also be useful in mapping new data. For example, although Horizontal cells and Retinal ganglion cells were mapped to the region adjacent to/overlap the region occupied by Amacrine cells, these cells exhibited low log-likelihoods. We therefore should not simply conclude that these cells were Amacrine cells, but need further analyses to elucidate these cell types.

`scvis` preserves the ‘global’ structure in a dataset, greatly enhancing interpretation of projected structures single-cell RNA-sequencing data. For example, in the bipolar dataset, we can see that the ‘ON’ bipolar cells were close to each other in the two-dimensional representation in Fig. 3(a), and similarly, the ‘OFF’ bipolar cells were close to each other. For the oligodendroglioma dataset, the cells can be first divided into normal cells and malignant cells, the normal cells formed two clusters, with each cluster of cells consisting of cells from multiple patients. The malignant cells, although from the same patient, formed multiple clusters with cell clusters from the same patient adjacent to each other. Adjacent malignant cell clusters from different patients tended to selectively express the oligodendrocyte markers genes or the astrocyte marker genes. For the metastatic melanoma dataset, malignant cells from different patients, although mapped to the same region, formed clusters based on the patient origin of the cells, while immune cells from different patients tended to be clustered together by cell types. From the low-dimensional representations, we can hypothesize that the cancer-associated fibroblasts were more ‘similar’ to the malignant cells than to the immune cells.

Other methods, e.g., the SIMLR algorithm, improve the t-SNE algorithm^49^ by learning a similarity matrix between cells, and the similarity matrix is used as the input of t-SNE for dimension reduction. However, SIMLR is computationally expensive because its objective function involves large matrix multiplications (an *N × N* kernel matrix multiplying an *N × N* similarity matrix, where N is the number of cells). In addition, although the learned similarity matrix could help clustering analyses, it may distort the manifold structure as demonstrated in the t-SNE plots on the learned similarity matrix^49^ because the SIMLR objective function encourages forming clusters. The most similar approach for scvis may be the parametric t-SNE algorithm^50^, which uses a neural network to learn a parametric mapping from the high-dimensional space to a low dimension. However, parametric t-SNE is not a probabilistic model, the learned low-dimensional embedding is difficult to interpret, and there is no likelihoods to quantify the uncertainty of each mapping.

In conclusion, the scvis algorithm provides a computational framework to compute low dimensional embeddings of single cell RNA-sequencing data while preserving global structure of the high dimensional measurements. We expect scvis to model and visualize structures in single-cell RNA-sequencing data while providing new means to biologically interpretable results. As technical advances to profile the transcriptomes of large numbers of single cells further mature, we envisage that scvis will be of great value for routine analysis of large-scale, high resolution mapping of cell populations.

## Methods

### A latent variable model of single-cell data

We assume that the gene expression vector **x***_n_* of cell n is a random vector and is governed by a low-dimensional latent vector **z**_*n*_. The graphical model representation of this latent variable model (with *N* cells) is shown in Fig. 5(a). The **x**_*n*_ distribution could be a complex high-dimensional distribution. We assume that it follows a Student’s *t*-distribution given **z**_*n*_:

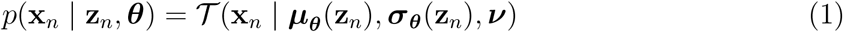

where both ***μ**_θ_*(.) and ***σ****_θ_* (.) are functions of **z** given by a neural network with parameter ***θ***, and *v* is the degree of freedom parameter and learned from data. The marginal distribution 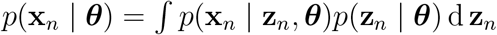 can model a complex high-dimensional distribution.

**Figure 5:**
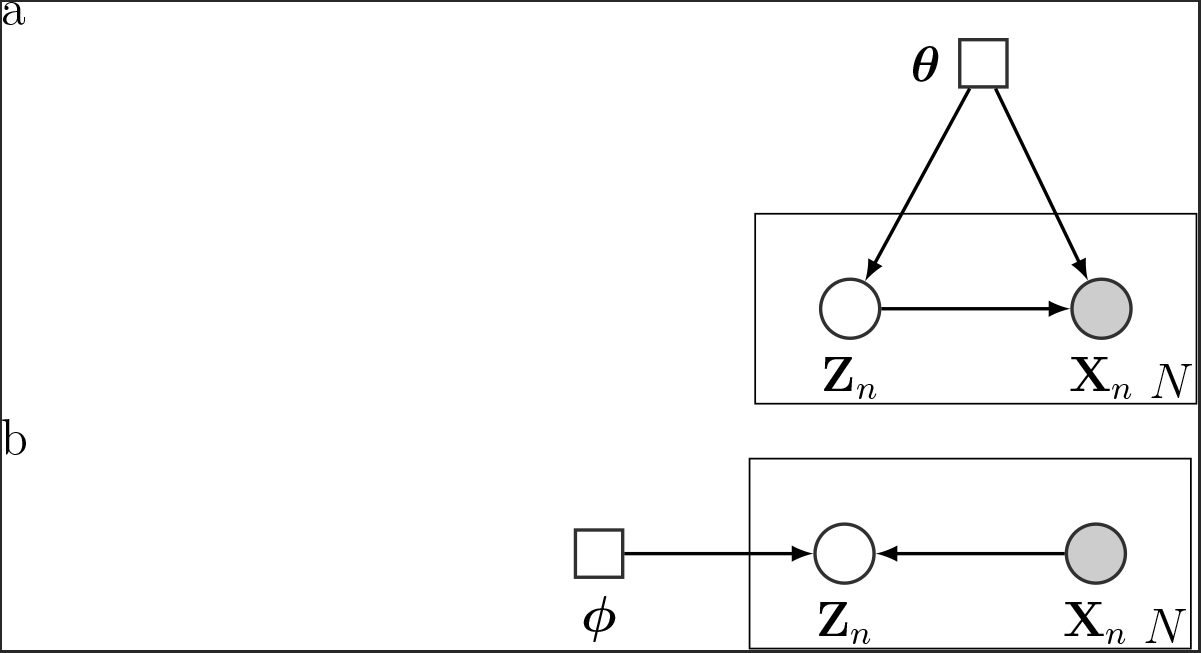
The scvis directed probabilistic graphical model and the variational approximation of its posterior. Circles represent random variables. Squares represent deterministic parameters. Shaded nodes are observed, and unshaded nodes are hidden. Here we use the plate notation, i.e., nodes inside each box will get repeated when the node is unpacked (the number of repeats is on the bottom right corner of each box). Each node and its parents constitute a family. Given the parents, a random variable is independent of the ancestors. Therefore, the joint distribution of all the random variables is the products of the family conditional distributions. (a) The generative model to generate data xn, and (b) the variational approximation *q*(**z***_n_* | **x***_n_*, *ϕ*) to the posterior *p*(**z**_*n*_ | **x**_*n*_, ***θ***).

We are interested in the posterior distribution of the low-dimensional latent variable given data: *p*(**z**_*n*_ | **x***_n_*, ***θ***), which is intractable to compute. To approximate the posterior, we use the variational distribution *q*(**z***_n_* | **x***_n_*, *ϕ*)=*N* (*μ_ϕ_* (**x***_n_*), diag(***σ****_ϕ_* (**x***_n_*))). Both ***μ**_ϕ_*(.) and ***σ**_ϕ_*(.) are functions of x through a neural network with parameter *ϕ*. Although the number of latent variables grows with the number of cells, these latent variables are governed by a neural network with a fixed set of parameters *ϕ*. Therefore, even for datasets with large number of cells we still can efficiently infer the posterior distributions of latent variables. The model coupled with the variational inference is called the variational autoencoder^51,52^.

Now the problem is to find the variational parameter *ϕ* such that the approximation *q*(**z***_n_* | **x***_n_*, *ϕ*) is as close as possible to the true posterior distribution *p*(**z***_n_* | **x***_n_*, ***θ***). The quality of the approximation is measured by the Kullback-Leibler 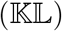 divergence^53^

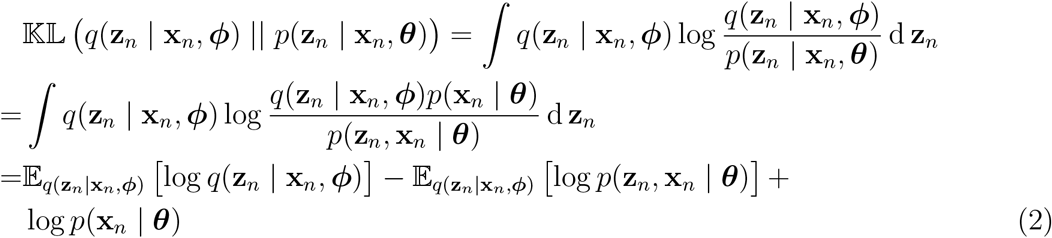

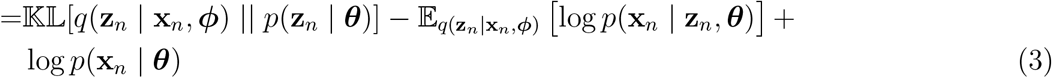

The term 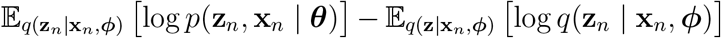 in Equation 2 is the evidence lower bound (ELBO) because it is a lower bound of log *p*(**x***_n_* | *θ*)as the 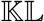 divergence on the left hand side is non-negative. We therefore can do maximum-likelihood estimation of both ***θ*** and *ϕ* by maximizing the ELBO. Notice that in the Bayesian setting, the ELBO is a lower bound of the evidence log *p*(**x**_*n*_) as the parameters ***θ*** are also latent random variables.

Both the prior *p*(**z**_*n*_ | ***θ***) and the variational distribution *q* (**z**_*n*_ | **x**_*n*_, ***ϕ***)intheELBO of the form in Equation 3 are distributions of **z**_*n*_. Inourcase, we can compute the 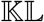 term analytically because the prior is a multivariate normal distribution, and the variational distribution is also a multivariate normal distribution g_×_iven xn. However, typically there is no closed-form expression for the integration 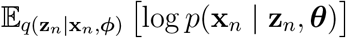 because we should integrate out **z**_*n*_ and the parameters of the model ***μ**_θ_* (**z**_*n*_) and diag(*σ_θ_* (**z***_n_*)) are functions of **z***_n_*. Instead, we can use Monte-Carlo integration and obtain the estimated evidence lower bound for the n-*th* cell:

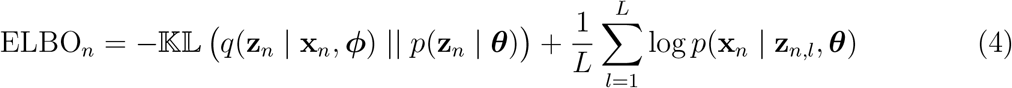

where z*_n,l_* is sampled from *q*(**z**_*n*_ | **x**_*n*_, *ϕ*)and *L* is the number of samples. We want to take the partial derivatives of the evidence lower bound w.r.t. the variational parameter *ϕ* and the generative model parameter ***θ*** to find a local maximum of the ELBO. However, if we directly sample points from *q* (**z***_n_* | x*n*, *ϕ*), it is impossible to use the chain rule to take the partial derivative of the second term of Equation 4 w.r.t *ϕ* because z*_n,l_* is a number. To use gradient based methods for optimization, we indirectly sample data from *q* (**z**_*n*_ | **x**_*n*_, *ϕ*) using the ‘reparameterization trick’^51,52^. Specifically, we first sample ϵ_l_ from a easy to sample distribution *ϵ_l_* ∼ *p*(*ϵ | **α***), e.g., a standard multivariate Gaussian distribution for our case. Next we pass ϵ*_l_* through a continuous funnction *g_ϕ_* (*ϵ, **x**_*n*_*) to get a sample from *q*(**z**_*n*_ | **x**_*n*_, *ϕ*). For our case if 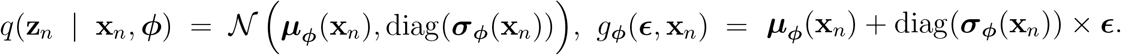

### Adding regularizers on the latent variables

Given *i.i.d* data 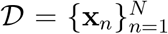, by maximizing the 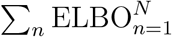, we can do maximum-likelihood estimation of the model parameters ***θ*** and the variational distribution parameters ***ϕ***. Although *p*(**z**_*n*_ | ***θ***)*p*(**x***_n_* | **z**_*n*_, ***θ***) may model the data distribution very well, the variational distribution *q* (**z***_n_* | **x***_n_*, *ϕ*) is not necessarily good for visualization purposes. Specifically, it is possible that there are no very clear gaps among the points from different clusters. In fact, to model the data distribution well, the low-dimensional **z** space tends to be filled such that all the **z** space is used in modeling the data distribution. To better visualize the manifold structure of a dataset, we need to add additional regularizers to the objective function in Equation 4 to encourage forming gaps between clusters, and at the same time keeping nearby points in the high-dimensional space nearby in the low-dimensional space. Here we use the t-distributed stochastic neighbour embedding (t-SNE)^34–39^ objective function.

The t-SNE algorithm preserves the local structure in the high-dimensional space after dimension reduction. To measure the ‘localness’ of a pairwise distance, for a data point *i* in the high-dimensional space, the pairwise distance between *i* and another data point *j* is transformed to a conditional distribution by centering an isotropic univariate Gaussian distribution at *i*

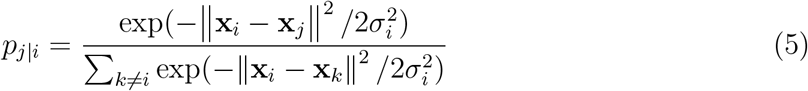

The point specific standard deviation *σ_i_* is a parameter which is computed automatically in such a way that the perplexity 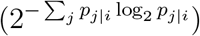 of the conditional distribution *p_j|i_* equals a user defined hyper-parameter (e.g., typically 30^54^). We set *p_i|i_* = 0 because only pairwise similarities are of interest.

In the low-dimensional space, the conditional distribution *q_j|i_* is defined similarly and *q_i|i_* is set to 0. The only difference is that an unscaled univariate Student’s *t*-distribution is used instead of an isotropic univariate Gaussian distribution as in the high-dimensional space. Because in the high-dimensional space, more points can be close to each other than in the low dimensional space (e.g., only two points can be mutually equidistant in a line, three points in a two dimensional plane, and four points in a three dimensional space), it’s impossible to faithfully preserve the high-dimensional pairwise distance information in the low-dimensional space if the intrinsic dimensionality of the data is bigger than that of the low dimensional space. A heavy tailed Student’s *t*-distribution allows moderate distances in the high-dimensional space to be modeled by much larger distances in the low dimensional space to prevent crushing different clusters together in the low-dimensional space^34^.

The low-dimensional embedding coordinates 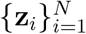 are obtained by minimizing the 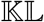 divergence between the sum of conditional distributions:

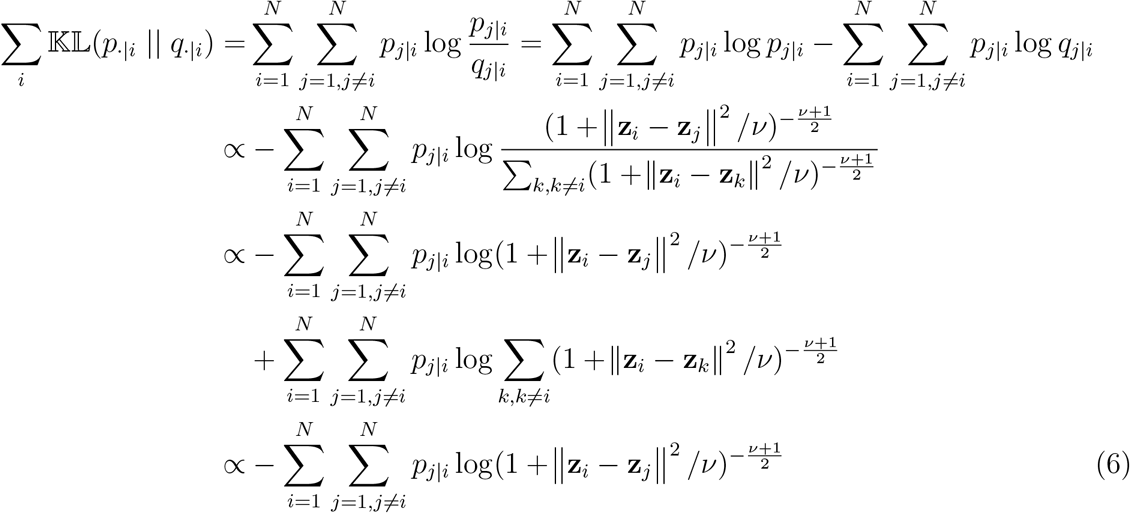

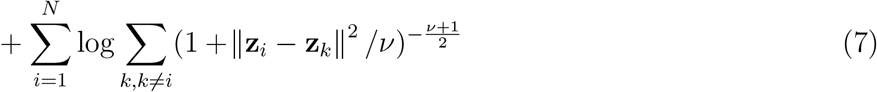

Here *v* is the degree of freedom of the Student’s t distribution, which is typically set to one (the standard Cauchy distribution) or learned from data. Equation 6 is a data dependent term (depending on the high-dimensional data) that keeps nearby data points in the highdimensional data nearby in the low-dimensional space^37^. Equation 7 is a data independent term that pushes data points in the low-dimensional space apart from each other. t-SNE has shown excellent results on many visualization tasks such as visualizing scRNA-seq data and CyTOF data^40^.

The final objective function is a weighted combination of the ELBO of the latent variable model and the t-SNE objective function:

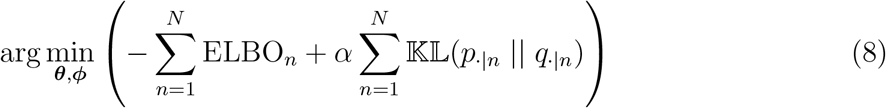

The parameter α is set to the dimensionality of the input high-dimensional data because the magnitude of the log-likelihood term in the ELBO scales with the dimensionality of the input data. The perplexity parameter is set to 10 for `scvis`.

### Datasets

The oligodendroglioma dataset measures the expression of 23,686 genes in 4,347 cells from six *IDH1* or *IDH2* mutant human oligodendrolioma patients^44^. The expression of each gene is quantified as log_2_ (TPM/10 + 1), where ‘TPM’ standards for ‘transcripts per million’^55^. Through copy number estimations from these scRNA-seq measurements, 303 cells without detectable copy number alterations were classified as normal cells. These normal cells can be further grouped into microglia and oligodendrocyte based on a set of marker genes they expressed. Two patients show sub-clonal copy number alterations.

The melanoma dataset is from sequencing 4, 645 cells isolated from 19 metastatic melanoma patients^3^. The cDNAs from each cell were sequenced by an Illumina NextSeq 500 instrument to 30bp pair-end reads with a median of ∼150,000 reads per cell. The expression of each gene (23,686 genes in total) is quantified by log_2_ (TPM/10 + 1). In addition to malignant cells, the authors also profiled immune cells, stromal cells, and endothelial cells to study the whole tumour multi-cellular ecosystem.

The bipolar dataset consists of low-coverage (median depth of 8,200 mapped reads per cell) Drop-seq sequencing^9^ of 27,499 mouse retinal bipolar neural cells from a transgenic mouse^1^. In total 26 putative cells types were identified by clustering the first 37 principal components of all the 27,499 cells. Fourteen clusters can be assigned to bipolar cells, and another major cluster is composed of Muller glia cells. These 15 clusters account for about 96% of all the 27,499 cells. The remaining 11 clusters (comprising of only 1,060 cells) include Rod photoreceptors, Cone photoreceptors, Amacrine cells, and cell doublets and contaminants^1^.

The retina dataset consists of low-coverage Drop-seq sequencing^9^ of 44,808 cells from the retinas of 14-day-old mice. By clustering the 2D t-SNE embedding using DBSCAN^56^ – a density-based clustering algorithm, the authors identified 39 clusters after merging the clusters without enough differentially expressed genes between any two clusters.

### Code and data availability

The `scvis` Python package will be made available from bit-bucket: https://bitbucket.org/jerry00/scvis-dev. All scRNA-seq data analyzed in this paper are publicly available from the single cell portal (https://portals.broadinstitute.org/single_cell), or from the Gene Expression Omnibus (bipolar: GSE81905, retina: GSE63473, oligodendroglioma: GSE70630, metastatic melanoma: GSE72056).

## Acknowledgements

This work was supported by a Discovery Frontiers project grant, “The Cancer Genome Collaboratory”, jointly sponsored by the Natural Sciences and Engineering Research Council (NSERC), Genome Canada (GC), the Canadian Institutes of Health Research (CIHR) and the Canada Foundation for Innovation (CFI) to S.P.S. In addition, we acknowledge generous long-term funding support from the BC Cancer Foundation. The S.P.S. group receives operating funds from the Canadian Breast Cancer Foundation, the Canadian Cancer Society Research Institute (impact grant 701584 to S.P.S.), the Terry Fox Research Institute (PPG program on forme fruste tumors), CIHR (grant MOP-115170 to S.P.S.), CIHR Foundation (grant FDN-143246 to S.P.S.). S.P.S. is supported by Canada Research Chairs. S.P.S. is a Michael Smith Foundation for Health Research scholar.

## Author’s contributions

J.D., project conception, software implementation, and data analysis; J.D., A.C., and S.P.S., algorithm development and manuscript writing; S.P.S., project conception, oversight, and senior responsible author.

